# Graspable foods and tools elicit similar responses in visual cortex

**DOI:** 10.1101/2024.02.20.581258

**Authors:** J. Brendan Ritchie, Spencer Andrews, Maryam Vaziri-Pashkam, Christopher I. Baker

## Abstract

Extrastriatal visual cortex is known to exhibit distinct response profiles to complex stimuli of varying ecological importance (e.g., faces, scenes, and tools). The dominant interpretation of these effects is that they reflect activation of distinct “category-selective” brain regions specialized to represent these and other stimulus categories. We sought to explore an alternative perspective: that the response to these stimuli is determined less by whether they form distinct categories, and more by their relevance to different forms of natural behavior. In this regard, food is an interesting test case, since it is primarily distinguished from other objects by its edibility, not its appearance, and there is evidence of food-selectivity in human visual cortex. Food is also associated with a common behavior, eating, and food consumption typically also involves the manipulation of food, often with the hands. In this context, food items share many properties in common with tools: they are graspable objects that we manipulate in self-directed and stereotyped forms of action. Thus, food items may be preferentially represented in extrastriatal visual cortex in part because of these shared affordance properties, rather than because they reflect a wholly distinct kind of category. We conducted fMRI and behavioral experiments to test this hypothesis. We found that behaviorally graspable food items and tools were judged to be similar in their action-related properties, and that the location, magnitude, and patterns of neural responses for images of graspable food items were similar in profile to the responses for tool stimuli. Our findings suggest that food-selectivity may reflect the behavioral affordances of food items rather than a distinct form of category-selectivity.

## Introduction

An important feature of extrastriatal visual cortex is that it exhibits distinct neural response profiles to complex stimuli of varying ecological importance, including faces, bodies, scenes, tools, words, and object forms more generally (1,2). The dominant interpretation of such findings is that they reflect activation of distinct brain regions specialized to represent these and other stimulus categories; in other words, they reflect forms of “category-selectivity” in visual cortex. However, an alternative perspective is that the neural response to these stimuli is determined less by whether they form distinct categories, and more by their relevance to different forms of natural behavior, such as social interaction, navigation, and object manipulation (1,3,4). We sought to provide evidence for this alternative view by focusing on food.

Although food is primarily distinguished from other objects and materials in our environment by its edibility, not its appearance, many lines of evidence suggest that regions of visual cortex respond differentially to images of food compared to other types of objects. For example, early studies on food-selectivity found that images of food activate different portions of ventral occipitotemporal cortex (OTC) relative to non-food stimuli (5,6). Studies on food and taste related properties have also found that information about food type, taste qualities, whether food is processed, and even taste-related mental imagery can be decoded from OTC (7–9). These findings have been further corroborated by three recent studies using data-driven approaches (10–12). Each of these studies separately analyzed the Natural Scenes Dataset (NSD), which consists of fMRI BOLD responses to thousands of complex natural scene images annotated for different objects, including many types of foods (13). In each case, these studies found that OTC differentially represented stimuli based on whether there was food present in the scene or not (14). In the present study we sought to determine how the neural response to food related stimuli should be situated within the functional organization of occipitotemporal cortex more broadly.

A natural interpretation of these findings is that they reflect an underappreciated form of category-selectivity: food items are another ecologically important stimulus that is preferentially represented by OTC. For example, the studies that reported evidence of food selectivity based on the analysis of the NSD found that food was a distinct component in explaining variation in the NSD alongside these other stimulus types that are associated with different category-selective brain regions (10–12). Selectivity for food has also been compared to forms of category-selectivity in earlier work as well (5,15). At the same time, food is obviously also associated with a common behavior: eating. Food consumption typically involves the manipulation of food, and when this is done directly with the hands, food items share many properties in common with tools: they are graspable objects that we manipulate in self-directed and stereotyped forms of action. Thus, food items may be preferentially represented in OTC in part because of these shared affordance properties, rather than because they reflect a wholly distinct kind of object category. Earlier studies in some cases also compared the neural responses of food items and tools, with tools even being used as a non-food stimulus contrast class to identify food-selective brain regions (6,15,16). However, these studies did not select food stimuli based on their relative graspability, and as they defined food selectivity as a greater response for food than tools, this would naturally omit areas commonly activated by both stimuli.

We conducted an fMRI experiment to determine whether the location, magnitude, and spatial patterns of BOLD responses to graspable food items is similar to that for tools. We designed a stimulus set that allowed for comparing the response to graspable food items and tools, which we validated using a number of behavioral measures. We compared the whole brain response to graspable food items and tools to determine whether they localized similar regions of occipitotemporal and occipitoparietal cortex. We then evaluated the magnitude and pattern of neural responses to graspable food items in tool-selective regions of interest (ROIs). In line with our hypothesis, we found that, graspable food items elicited a similar response profile to tools in visual cortex.

## Results

### Behavioral evidence that graspable food items are represented similarly to tools

The primary stimulus set consisted of images of foods, tools, manipulable objects, and animals (**Fig 1A**). To control for shape, the objects were selected to exhibit similar variation in aspect-ratio and orientation across object types (17). To validate that the food items were represented as having similar properties to tools, several behavioral experiments were carried out online by a large number of participants who were separate from the fMRI participant pool. These tasks were designed to determine what properties differentiate tools from other types of manipulable objects and also assess the relative similarity of tools based on how they are manipulated during action.

**Fig 1.**
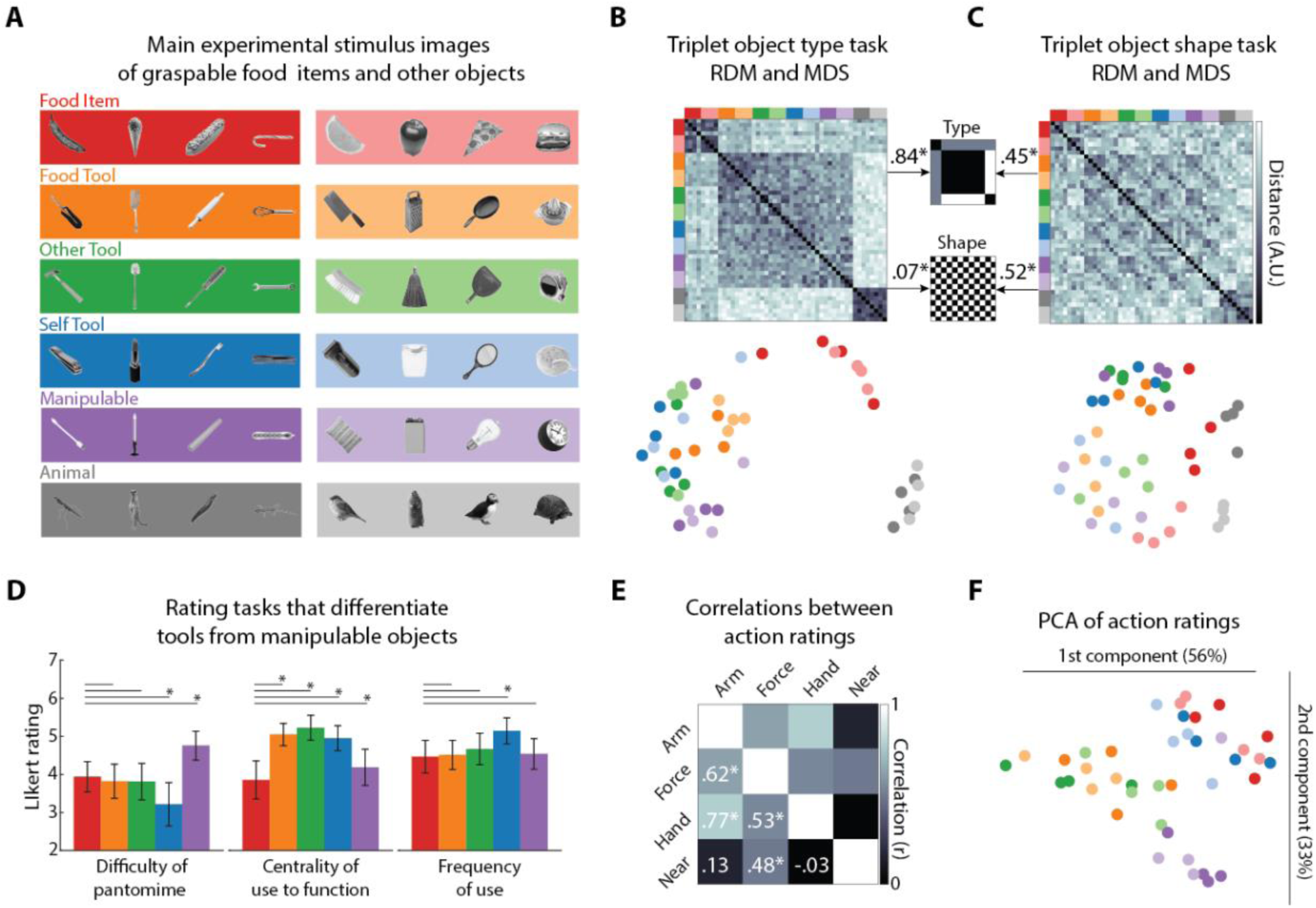
Primary stimuli and behavioral results. (A) The 48 object images used in the behavioral tasks and event-related fMRI runs. Images consisted of 8 exemplars each of six object types: Food Items, Food Tools, Other Tools, Self Tools, Manipulable objects, and Animals. Exemplars have two broad types of aspect ratios (darker color = higher aspect ratio; lighter color = lower aspect ratio), and had variable, matched orientation across object types. (B) Group-averaged RDM and 2D MDS solution for the object type triplet task. (C) Group-averaged RDM and 2D MDS solution for the object shape triplet task. For (B) an (C) Spearman’s ρ correlations between triplet task and model RDMs. * = p < 0.05. (D) bar plots of the mean Likert ratings for the three tasks used to differentiate tools from manipulable objects. * = p < 0.05. Error bars are the 95% CI for a normal distribution centered on the mean. (E) Correlation matrix for the mean ratings for the four action rating tasks. * = p < 0.05. (F) First two dimensions from PCA applied to the mean ratings of the four action rating tasks.

Two separate odd-one-out similarity tasks were performed by different groups of participants who judged which object in a triplet of objects was more distinct from the other two (18). These judgments were used to construct representational dissimilarity matrices (RDMs) of all stimulus pairs (**Fig 1B-C**). In the object type task, participants judged object similarity while ignoring shape. The mean split-half reliability of the RDM for this task was high (*r* = 0.86). To quantify the structure of the object type RDM, we correlated it with a model matrix that treated food items as intermediate in their similarity relative to the tools and manipulable objects compared to the animal stimuli and a binary shape matrix based on the aspect ratio of the objects. These models were not correlated with each other (ρ = -0.02, p = 0.49). We found that the object type RDM was highly correlated with the intermediate similarity model (ρ = 0.84, p = 6.51e-294), and was also weakly correlated with the binary shape model (ρ = 0.07, p = 0.02). That the structure of the RDM is well-captured by the intermediate similarity model can be seen in the results of the 2D multidimensional scaling (MDS) analysis that shows food items as more similar to artefacts than animals. In the 2D space artefacts (tools and manipulable objects) cluster together and are separate from animals (**Fig 1B**). Thus, the graspable food items we selected were judged as more similar to tools compared to animals.

In the Object Shape task, a separate set of participants judged object shape similarity while ignoring object type. The mean split-half reliability of the RDM for this task was high (*r* = 0.73). This RDM was found to be robustly correlated with the binary shape model (ρ = 0.52, p = 1.46e-77), and also, surprisingly, the intermediate similarity model (ρ = 0.45, p = 2.77e-58). This latter result suggests that although participants were instructed to ignore object type, this was nonetheless reflected in fine-grained comparisons. The two components of the RDM can also be seen in the MDS solution (**Fig 1C**). Food and animals clustered somewhat separated from the other artefact stimuli, which may reflect the fact that, even when ignoring object type, these objects are grouped together because of similarities in their shape properties (e.g., animals having limbs). As would be expected from the preceding findings, the object type and shape RDMs were also correlated with each other (ρ = 0.57, p = 5.37e-98), a result we return to in later analysis.

We also asked separate groups of participants to rate the food and artefact stimuli based on three tasks, adapted from Mahon et al. (19). These tasks were developed to differentiate between tools and other equally manipulable objects. For tools, there is a systematic relationship between form and how they are manipulated to serve a function, while for other equally manipulable objects there is no such relationship (**Fig 1D**). For each task, the individual stimulus responses were reliable across participants (mean split-half reliabilities: Difficulty of Pantomime, *r* = 0.85; Centrality of Use to Function; *r* = 0.85; Frequency of Use, *r =* 0.75). We examined whether the mean ratings of difficulty for the Food Items were significantly different from those for the different types of artefacts. For the difficulty of Pantomime task, the mean ratings were significantly greater than that for the Self Tools (t(40) = 3.61, p = 8.54e-04) and lower than that for the Manipulable Objects (t(40) = -3.1, p = 0.004). In line with the findings of Mahon and colleagues, participants on average found it more difficult to imagine pantomiming use of Manipulable Objects, while Food Items were comparable to most tools. For the Centrality of Use task, the mean ratings were significantly lower for Food Items than all other objects, even compared to the Manipulable Objects (t(39) = -4.1, p = 2.13e-04). This is presumably because the function of food is to provide sustenance, which is not overtly tied to how it is manipulated. Finally, for the Frequency of Use task, the mean responses for the Food Items were only significantly lower than those for the Self Tools (t(38) = -2.9, p = 0.006), which were objects commonly used in self-care. Thus, graspable food items share properties with tools that differentiate them from manipulable objects and showed similar action properties to self-directed tools.

We also asked additional groups of participants to rate the actions of using the non-animal objects in four tasks adapted from Watson & Buxbaum (20), which were developed to characterize the action similarity between different tools. The responses for the stimuli were reliable across participants (mean split-half reliabilities: Arm Movement, *r =* 0.75; Force, *r* = 0.93; Hand Contact, *r* = 0.84; Near Body, *r* = 0.94). The mean ratings for the Arm Movement task were significantly correlated with those for Force (*r* = 0.62, p = 2.54e-05) and Hand Contact (*r* = 0.77, p = 5.35e-09) tasks, and Force and Hand Contact task responses were also significantly correlated (*r* = 0.53, p. = 4.98e-04). Of the other tasks, only the Force responses were also correlated those for the Near Body task (*r* = 0.48, p = 0.002). Following (20), we used principal component analysis (PCA) to characterize a 2D space, which captured most of the variance in the mean ratings for all four tasks (89%). In this 2D space the Food Items and Self Tools clustered separately from the other types of tools along the first component (**Fig 1F**). An unexpected result of this analysis was that Manipulable Objects also clustered separately from the food and tool stimuli on the second component.

In summary, these behavioral results provide evidence that cognitively, graspable food items are represented as overall more similar to tools than animals and have many properties comparable to tools and manipulable objects.

### Graspable food items and tools elicit similar loci of activity in visual cortex

Tool-selective regions of visual cortex have been observed in anterior left lateral (LOTC) and medial bilateral (LOTC) regions of OTC, and in many regions within or close to the post-central sulcus (PCS) and supra marginal gyrus (SMG) in parietal cortex (21). These regions have typically been isolated using functional localizers that contrast the evoked response of tools > animals (22–25). We sought to determine whether a functional contrast of graspable food items > animals would identify similar or overlapping regions of activity in visual cortex, and also compared the results of an objects > animals contrast (**Fig 2A ‘)**. These contrasts were tested at the group level with mixed effects meta-analysis using the normalized beta weight and t statistic maps of the three contrasts of interest (26).

**Fig 2.**
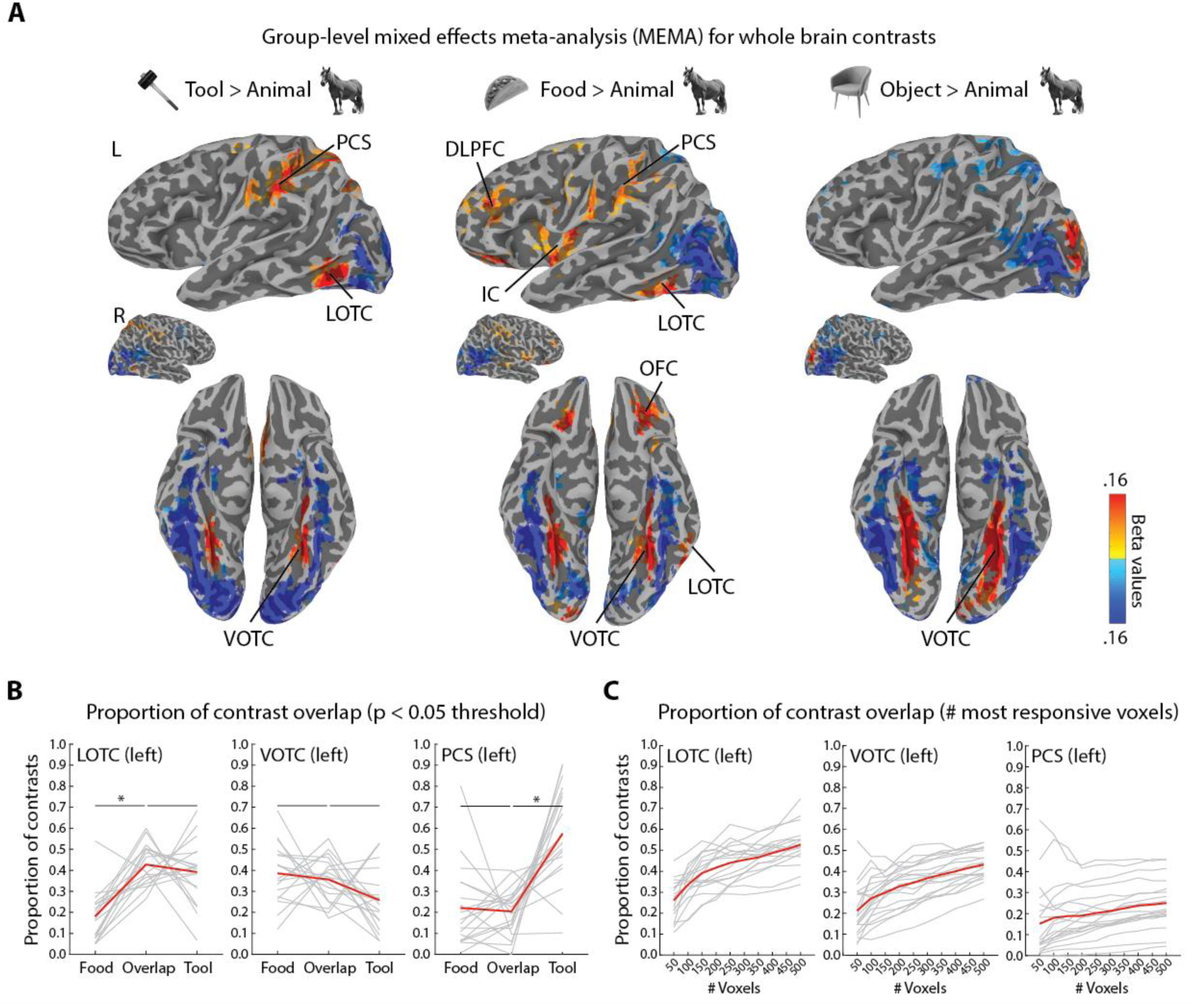
Overlap between tool- and food-selective regions of visual cortex. (A) Group level contrast results analyzed using mixed effects meta-analysis (MEMA) (Chen et al. 2012). All results are thresholded at t(19) = 2.09, p < 0.05, two-sided, uncorrected. Voxels cluster threshold = 40. DLPFC = dorsolateral prefrontal cortex; IC = insular cortex; LOTC = lateral occipitotemporal cortex; OFC = orbitofrontal cortex; PCS = post-central sulcus; and VOTC = ventral occipitotemporal cortex. (B) Proportion of overlap between loci of activation for food and tool selectivity in LOTC, VOTC, and PCS. Individual contrasts were thresholded at p < 0.05 and proportion of overlap reflects the union of the resulting regions of selectivity (i.e, the Jaccard Index). * = p < 0.05. (C) Proportion of overlap (Jaccard Index) based on the increasing numbers of most selective voxels in LOTC, VOTC, and PCS.

Three contrasts were carried out (**Fig 2A**). First, the standard tool > animal contrast revealed three group-level loci of activation: (i) left lateral anterior LOTC, (ii) bilateral medial VOTC, and (iii) predominantly left lateralized PCS. These results are analogous to numerous studies on tool-selectivity in visual cortex (21,27). Second, the food > animal contrast revealed a superficially very similar trio of activation regions in visual cortex as the tool > animal contrast, along with bilateral activation in insular cortex (IC), dorsolateral prefrontal cortex (DLPFC), and orbitofrontal cortex (OFC). These latter findings also replicate the results of prior studies on the response to food images (5,7). Finally, an object > animal contrast did not reveal activation in either LOTC or PCS, but it did produce a strong bilateral response in VOTC. Thus, graspable food items, unlike other objects, do produce similar loci of activation in visual cortex as tools.

We next calculated the amount of overlap in activation in left LOTC, VOTC, and PCS at the individual level, for both the tool > animal and food > animal contrasts (**Fig 2B**). This was done in two ways in participants’ native brain spaces using the Jaccard index, or the intersection of the maps above a threshold divided by their union (28). First, we selected all voxels within anatomical masks of these three swaths of cortex at a liberal statistical threshold of p < 0.05 and then determined the proportions of the intersection of the tool- and food-selective patches overlapped (the Jaccard index), or what portion of these maps did not overlap and so were unique tool- or food-selective responses. This operation could be carried out for the majority of participants in each mask region, since only a few participants did not show activation above threshold for both contrasts (LOTC-tool: N = 18; VOTC: N = 18; and PCS: N = 19). For LOTC, the overlap was on average ∼ 43 %, and the mean was significantly greater than for the food-unique patch (t(17) = - 6.6, p = 4.275e-06), but not the tool-unique patch ((t(17) = 0.68, p = 0.51). Thus, in LOTC most of the food-selective patches tended to consist of overlap with the tool-selective patches, which also appeared relatively stable across participants (**Fig 2B**). For VOTC, the overlap was on average ∼ 36 %, and was not significantly greater than that of the food-unique (t(17) = 0.61, p = 0.55) or tool-unique patch ((t(17) =1.93, p = 0.07). Thus, in VOTC the overlap area was on average comparable to the proportion of non-overlapping patches, though this may reflect heterogeneity at the individual level (**Fig 2B**). For PCS, the overlap on averages was ∼ 20 % and was not significantly greater than mean size of the food-unique patch (t(17) = 0.34, p = 0.74), and was significantly smaller than the tool-unique patch ((t(17) = -5.3, p = 4.88e-05). Thus, in PCS the overlap was on average comparable to the food-unique patch, but a large portion of the tool patch did not overlap, which was relatively consistent across participants (**Fig 2B**).

The above method of calculating overlap does not tell us however whether the overlap in activation is more at the peaks or the margins of patches of activity. Thus, next we determined the overlap by calculating the Jaccard index amongst the top 50 to 500 most responsive voxels for the two contrasts, in steps of 50 voxels (**Fig 2C**). For left LOTC and VOTC there were relatively steady increases in overlap as the size of voxel bin increased from ∼ 20 % upwards (to ∼ 50 % and ∼ 40 %, respectively), while for PCS even at 500 voxels PCS overlap remained < 30 %. Thus, at the individual level there was considerable overlap in patches of activity for food and tool contrasts, especially in LOTC and VOTC. Taken together, these results provide evidence that the spatial distribution of evoked neural responses to graspable food items in visual cortex are very similar to those for tools, when assessed based on the sorts of functional contrasts used to identify tool-selective portions of visual cortex.

### Similar responses for food items and tools in lateral and ventral OTC

Within classically tool-selective regions, tool images produce greater magnitude neural responses, and distinct patterns of activity, relative to non-tool stimuli such as animals. We next investigated whether this might also be the case for the Food Item stimuli in three tool-selective regions of interest (ROIs) in the left hemisphere, which could be defined from the tool > animal contrasts for most participants: LOTC-tool (N = 19), VOTC-tool (N = 18), and PCS-tool (N = 20). For comparison, we measured the response to food items in two ROIs defined based on the reverse contrast, animal > tool: LOTC-animal and VOTC-animal. We also tested whether some of the behavioral measures used to validate the food item stimuli might correlate with the neural responses in the tool-selective ROIs.

First, we assessed whether on average graspable Food Items produced similar magnitudes of response in the ROIs when compared to the different types of tools as well as manipulable object and animal stimuli. In the tool-selective ROIs, Food Items produced qualitatively comparable responses to the different tool and artefact exemplars relative to the Animal stimuli (**Fig 3A**). Averaging the responses for individual images, Food Items elicited significantly greater activation than Animals in all three tool-selective ROIs (**Fig 3A**): LOTC-tool (t(18) = 4.02, p = 8.05e-04), VOTC-tool (t(17) = 7.24, p = 1.39e-06), and PCS-tool (t(19) = 3.2, p = 0.005). The mean response was also significantly greater for food items in PCS-tool than for the Manipulable Objects (t(19) = 2.13, p = 0.046). In LOTC- and VOTC-tool, Food Items also produced on average lower responses than at least some of the tool and artefact stimuli. In LOTC-tool, the responses were significantly lower than for Other Tools (t(18) = -3.2, p = 0.005); and in VOTC-tool, they were significantly lower than for all tools and artefacts (t(17) ≤ -3.06, p ≤ 0.007). In the animal-selective ROIs, Food Items produced much lower responses than the Animal exemplars (**Fig 3A**). When averaging responses across exemplars, Food Items produced significantly lower activation in LOTC-animal than any other object type (t(19) ≤ -2.61, p ≤ 0.01). In VOTC-animal, mean responses were significantly lower than for animals (t(19) = -8.9, p = 3.27e-08), and greater than for Food Tools (t(19) = 3.07, p = 0.006).

**Fig 3.**
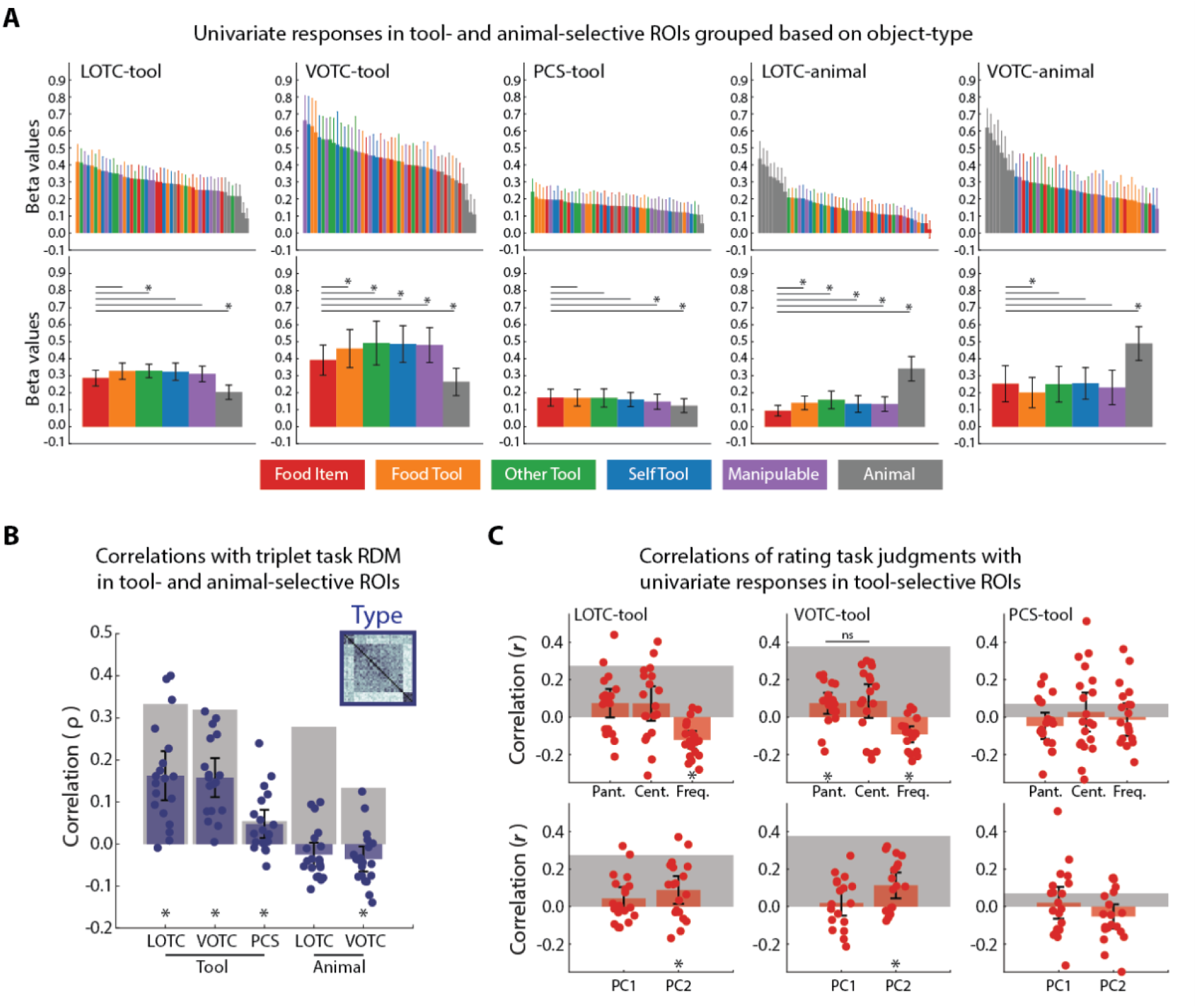
Univariate and multivariate responses in tool- and animal-selective ROIs in visual cortex. (A) Top row, mean univariate responses for individual images color-coded by object type in tool- and animal-selective ROIs. Bottom row, mean univariate responses in the ROIs after averaging across all exemplars of an object type. (B) Correlations between the object type RDM and neural RDMs for each of the ROIs. Grey bars indicate the noise ceiling of reliability for each ROI. Dots are data points for individual subjects (C) Top row, mean correlations between the mean ratings for the tasks from Fig 1D and the individual univariate responses for each of the tool-selective ROIs. Bottom row, mean correlations between the ratings for the two PCA dimensions from Fig 1F and individual univariate responses for each of the tool-selective ROIs. For both rows, grey block indicates the noise ceiling and dots are data points for individual subjects. For all plots, * = p < 0.05 (ns = not significant) and error bars are the 95% CI of a normal distribution centered on the mean.

Next, we constructed individual neural RDMs for each of the tool- and animal-selective ROIs, which we then correlated with the RDM for the object type triplet task. The mean correlations between RDMs were significant for each of the three tool-selective ROIs (t(17-19) ≥ 2.99, p ≤ 0.008), and were significantly negative for the animal-VOTC (t(19) = -2.49, p = 0.02). Thus, although the reliability of the animal-selective ROIs was similar to the tool-selective ROIs, their patterns of activity did not match the structure of the object type RDM. This result suggests that food items are also represented as intermediary in similarity between tools and animals in the pattern of responses in the tool-selective ROIs, which is also implied qualitatively when inspecting the group-averaged RDMs and MDS plots for these areas (**Fig 4A**) and comparing them to those for the animal-selective ROIs (**Fig 4B**).

**Fig 4.**
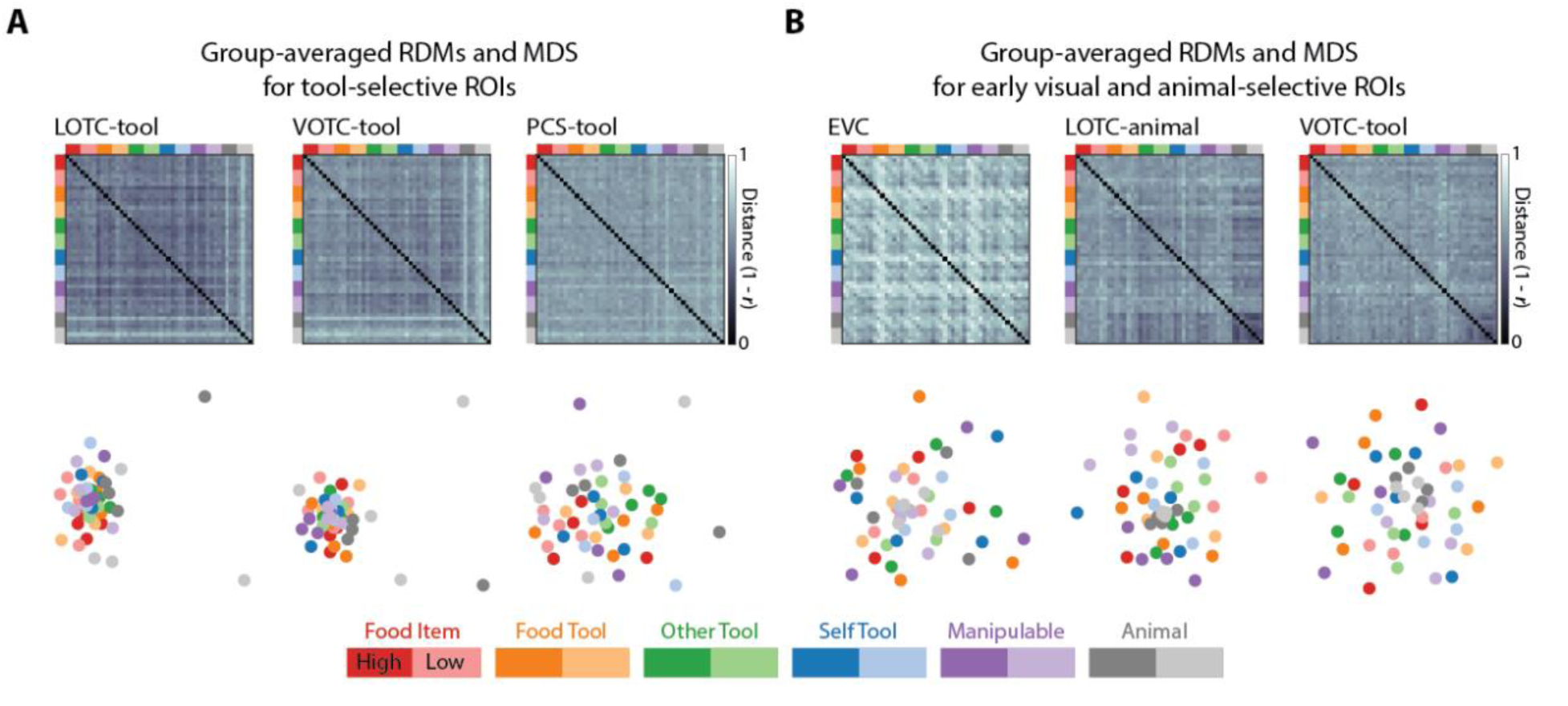
Representational geometry of tool- and animal-selective ROIs in visual cortex. (A) Group-averaged RDMs and 2D MDS solutions for the three tool-selective ROIs. (B) Group-averaged RDMs and 2D MDS solutions for the early visual cortex and animal-selective ROIs. Color saturation in legend indicates whether the stimuli were grouped as having high or low aspect ratio.

To determine whether the rating task judgments might predict some of the variance in the response magnitude in tool-selective ROIs, we also correlated the mean ratings with the magnitudes of responses across individual stimuli. For the three rating tasks for differentiating tools from Manipulable Objects (**Fig 3C**, top row), Frequency of Use was negatively correlated with responses in both LOTC-tool (t(18) = -5.46, p = 3.44e-05) and VOTC-tool (t(17) = -4.5, p = 3.16), which suggests that the strength of response is not positively related to how often an object is used. A significant effect was also found in VOTC-tool for the Pantomime ratings (t(17) = 2.77, p = 0.013), which was not significantly different from the correlation for the Centrality of Use ratings (t(17) = -0.19, p = 0.85). With respect to the action ratings (**Fig 3C**, bottom row), we correlated the first two principle components with the magnitude of responses in the tool-selective ROIs and found significant effects for PC2 in both LOTC-tool (t(18) = 2.5, p = 0.023) and VOTC-tool (t(17) = 3.46, p = 0.003). These results suggest that some of the semantic properties regarding the manipulation and use of the food and artefact stimuli may be represented in tool-selective regions of OTC. No effects were significant for PCS-tool.

In summary, the magnitude and pattern of response in tool-selective ROIs for food items at the individual level were greater and more similar to tools than animals. Furthermore, conceptual properties related to object manipulation for the stimuli may be encoded in tool-selective OTC.

### The representation of object shape type in tool-selective OTC and early visual cortex

It has been questioned whether effects of tool-selectivity in the visual cortex may result from biases in stimulus selection, since many canonical tools (e.g., a hammer) are elongated, or have higher aspect-ratio shape profiles, compared to other types of objects (29,30). Evidence for such a shape bias however is scant. Studies that have explicitly tested or controlled the shape properties across stimuli have found that the functionality of tools, rather than their shape, is a distinct driver of neural responses, especially in dorsal tool-selective cortex (27,31). Our stimuli were designed to control for the shape profiles of exemplars across object types (17). Still, we sought to test whether shape might modulate the magnitude and patterns of neural responses in tool-selective ROIs. Amongst tool-selective ROIs, we found that the mean response was greater for high- vs low-aspect ratio stimuli only in LOTC-tool, which contrasts with previous findings on tool-selective brain regions. Though qualitatively in LOTC-tool the magnitude of response, when stimuli were grouped based on aspect ratio, was still highly variable (**Fig 5A**). For comparison, we also carried out the same analysis for left early visual cortex (EVC) and found the reverse effect (t(19) = -6.18, p = 6.1e-9), likely reflecting the fact that a larger portion of the visual field is activated by the low-aspect ratio stimuli. To determine whether more fine-grained shape differences might be reflected in the patterns of activity in the ROIs, we compared the shape type RDM to the neural RDMs. Significant effects were observed for all three tool-selective ROIs (t(17–19) ≥ 2.98, p ≤ 0.008), but not EVC. Taken as a whole, these results suggest that aspect-ratio information is only reflected in the magnitude of responses in LOTC-tool, while more fine-grained shape differences may be reflected in the response patterns across tool-selective ROIs.

**Fig 5.**
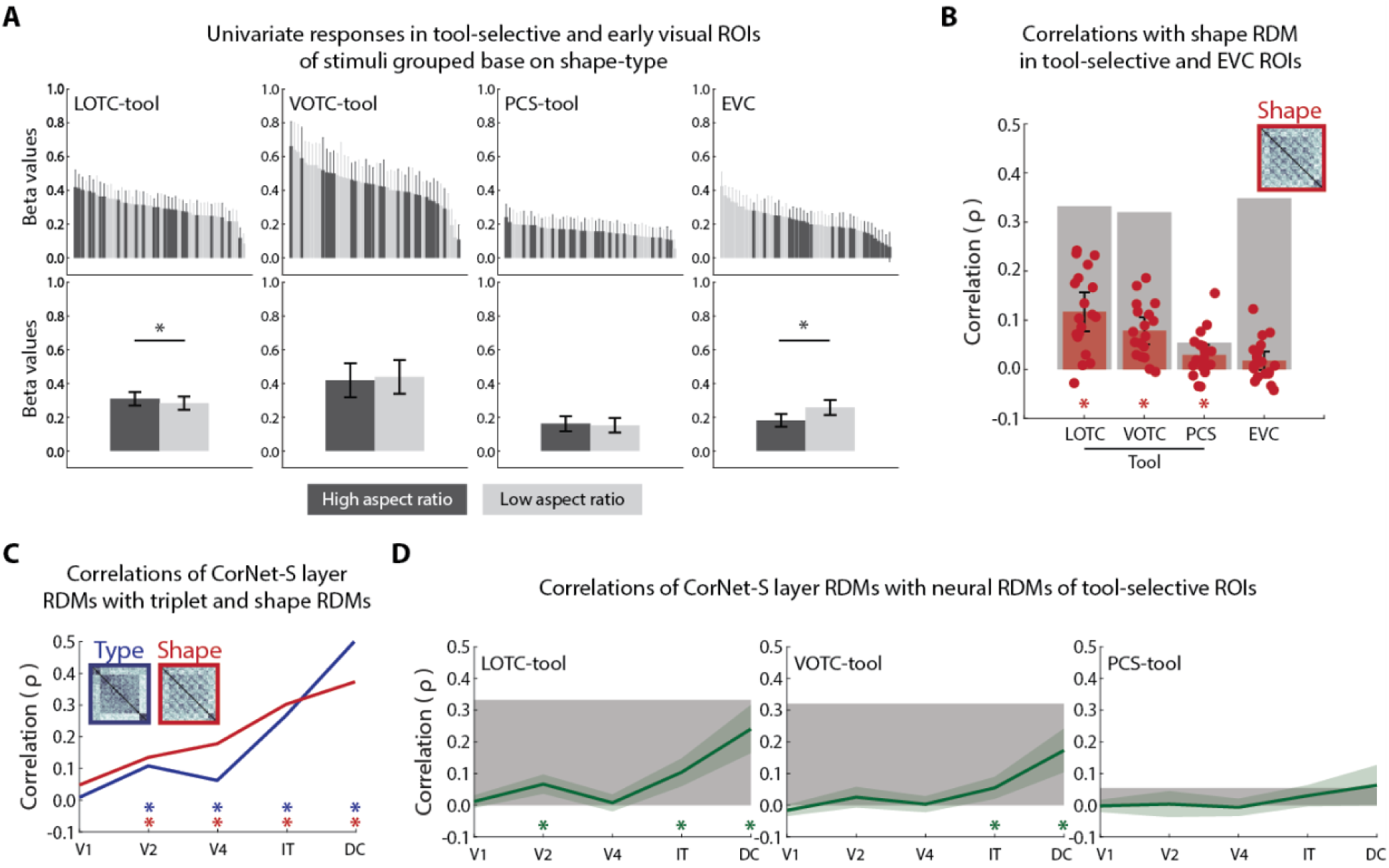
Represent of shape in tool-selective and early visual ROIs in visual cortex and comparisons of models and neural responses to a DNN. (A) Top row, mean univariate responses for individual images color-coded by shape type in tool-selective and early visual ROIs. Bottom row, mean univariate responses in the ROIs after averaging across all exemplars of a shape type. (B) Correlations between the object shape RDM and neural RDMs for each of the ROIs. Grey bars indicate the noise ceiling of reliability for each ROI. Dots are data points for individual subjects. (C) Correlations between triplet task RDMs and layers of CORnet-S. (D) Correlations between layer RDMs and neural RDMs for the three tool-selective ROIs. For all plots, * = p < 0.05 and error bars are the 95% CI of a normal distribution centered on the mean.

### Comparing the representation of food items and tools in a deep neural network

DNNs are a popular proxy model of visual processing for image labeling (32,33). Standardly trained DNNs learn to label images into different classes, including the object types targeted in the present study, like classes of tools, foods, and animals. Thus, one possibility is that they would have a similar representational structure to that reflected in the results of the object type triplet task, and the neural responses of tool-selective ROIs. This would suggest that the effects we have observed may not be driven by representations of functional and use properties of these objects (34). Thus, we sought to compare the layer representations of a DNN to the object type triplet task RDMs, to determine whether they contain a similar representation. We also compared the layer RDMs to the object shape RDMs, since object shape has been found to be represented separate from object type in the layers of DNNs (35). We then compared the layer RDMs for the DNN to the neural RDMs for the tool-selective ROIs. For this purpose we chose CORnet, a relatively shallow DNN designed to mimic the primate ventral visual stream (36). This DNN scores relatively high in the representational similarity between its IT layer and primate OTC on one common benchmark (37). The object and shape type RDMs were significantly correlated with the RDMs obtained from all but the first layer of CORnet (0.06 ≤ ρ ≤ .5, p ≤ 0.034), with highest correlations observed at the decoding (DC) layer (**Fig 5C**). When layer RDMs were correlated with the fMRI RDMs for the tool-selective ROIs, there were significant effects in the V2-, IT-, and DC-layers for LOTC-tool (t(18) ≥ 4.62, p ≤ 2.12e-04), and the IT- and DC-layers for VOTC-tool (t(17) ≥ 3.34, p ≤ 0.004). No significant correlations were observed between layer RDMs and fMRI RDMs for PCS-tool. These results show that a standardly trained DNN shows some representational similarity to not just the triplet task RDMs but also tool-selective regions of OTC.

### Partitioning the variance of BOLD dissimilarities in tool-selective visual cortex

Both the object type and shape triplet task RDMs were found to be correlated with multiple layers of CORnet. As mentioned earlier, the two triplet task RDMs were also found to be strongly correlated with each other. In light of this we sought to determine how much unique variance was accounted for by each of these models (34,38). Of particular interest was whether the relationships reflected in the triplet task RDMs are distinct from those found in the later layers of CORnet, since the former reflects how participants naturally grouped the objects while the latter was based solely on class labeling from images properties. When replotted and compared directly (**Fig 6A**), in LOTC-tool the mean correlations with the object type RDM were significantly greater than that with the object shape RDM (t(18) = 2.74, p = 0.013) and the RDM obtained from the IT-layer of CORnet (t(18) = 2.87, p = 0.01), while there was no significant difference between mean correlations with the object shape and IT-layer RDMs (t(18) =0.84, p = 0.41). The same pattern was observed for VOTC-tool (object type vs shape: t(17) = 5.91, p = 1.72e-05; object type vs IT-layer: t(17) = 6.93, p = 2.45e-06; object shape vs IT-layer: t(17) = 1.87, p = 0.08). There were no significant differences between the mean correlations for the three models in PCS-tool. We carried out variance partitioning by including all three models in multiple regression models with the group averaged neural RDMs as predictors (**Fig 6B**). In each case they significantly predicted a portion of the variance (0.04 ≤ *R*^2^ ≤ 0.15, p = 0). Furthermore, in each case the object type RDM by far accounted for the largest portion of unique variance amongst the three models. Notably, in contrast the shape type RDM accounted for almost no unique variance. These results suggest that the similarity relations reflected in the object type RDM, which, map onto the fMRI dissimilarity structures in tool-selective ROIs cannot be accounted for by a representation of a DNN that has learned to label stimuli as tools, food, and animals.

**Fig 6.**
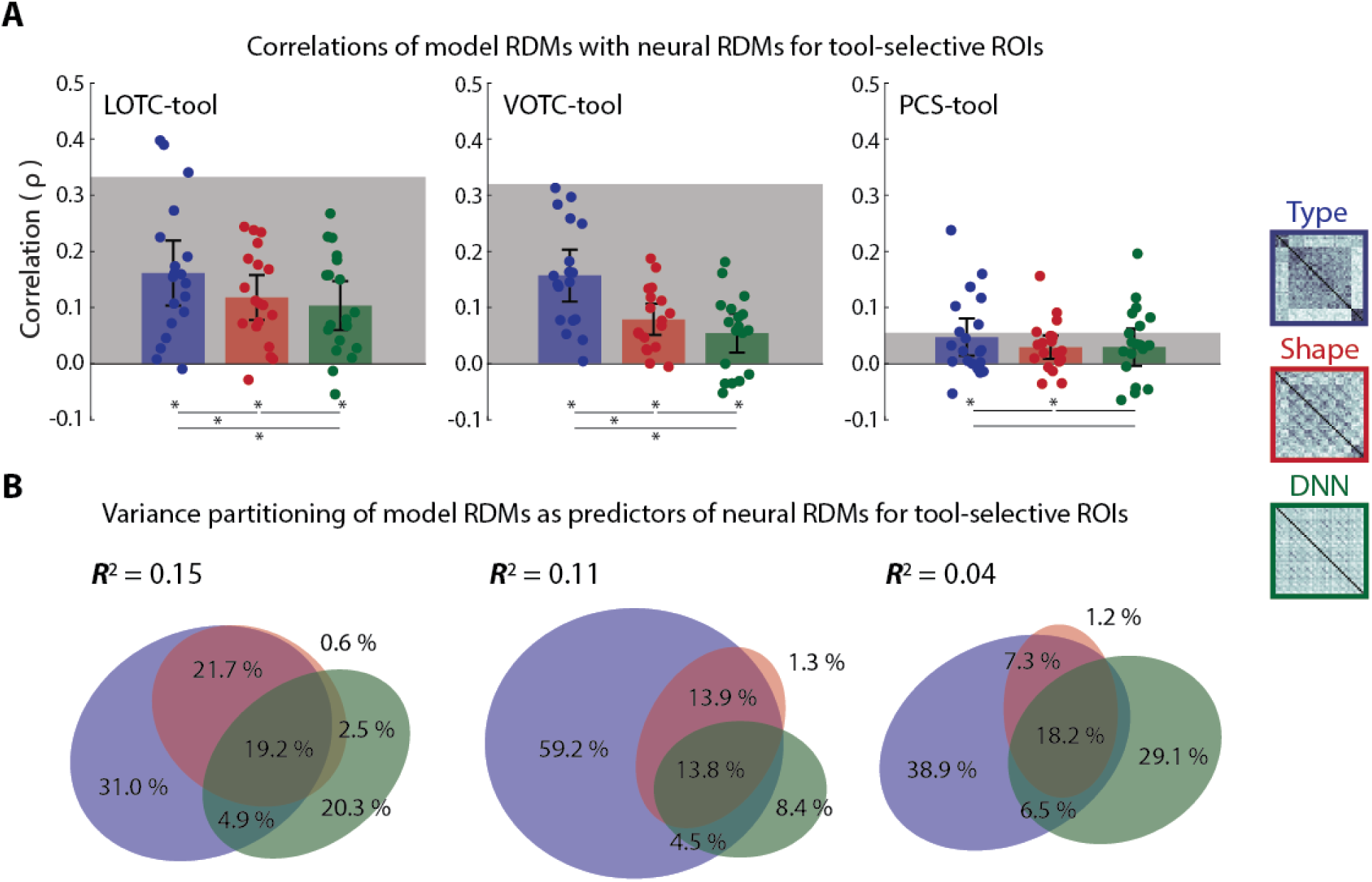
Variance partitioning for tool-selective ROIs in visual cortex. (A) Mean correlations of three model RDMs (object type triplet, object shape triplet, and IT-Layer of CORnet) with neural RDMs for the three tool-selective ROIs. * = p < 0.05 and error bars are the 95% CI of a normal distribution centered on the mean. (B) Results of variance partitioning (commonality analysis) for the three tool-selective ROIs. Coefficients of determination (*R*^2^) indicate the total proportion of explained variance for the full model. Regions of the Euler plots indicate percentages of the explained, and not total, variance accounted for by each component.

## Discussion

We tested the hypothesis that neural responses to images of graspable food items in the visual cortex may be similar in profile to those for tools since food items are also objects in our environment that are targets for manual manipulation and self-direct action. Based on behavioral judgements, we found that graspable food items are represented as more similar to tools than animals and share many properties related to action with manipulable objects. In an fMRI experiment we found that contrasting food and animal responses elicited very similar loci of neural activity in OTC and parietal cortex to tools as well as similar magnitudes and patterns of neural activity in tool-selective ROIs when compared to the neural response to animal stimuli. Looking at patterns of responses, the triplet object type model predicted a distinct amount of the variance when compared to the triplet object shape model and the layer representation of a DNN trained to represent label images of tool, food, and animal classes. Taken together, these results provide strong support for our hypothesis that the visual system similarly represents food and tools as targets of manipulable action. Our results have implications for: (i) whether visual cortex is specialized to represent food items as such, (ii) the characterization of “tool-selective” activity in visual cortex; (iii) the relationship between representations of object type and shape in visual cortex; and (iv) also point to a number of directions for future research.

### Making sense of food-selectivity in visual cortex

Some of the earliest studies providing evidence of food-selectivity in visual cortex used similar functional contrasts to those used to define regions of activity for stimuli like faces and scenes (5,16). A natural interpretation then of responses to food images is that since food items are of major ecological importance they are also represented as a privileged class of stimuli in the visual cortex. This interpretation has been further bolstered by recent studies using data-driven approaches to identify food-selectivity based on responses to a large number of complex natural scenes (10–12). In contrast, our results support the hypothesis that food responses in the visual cortex reflect food items as a kind of graspable object that we engage with during stereotyped forms of self-directed action. How then should these recent data-driven results and our own be reconciled?

It is notable that the three recent data-driven studies utilizing the NSD do not report any form of tool-selectivity although several types of tools are annotated in the stimulus set (e.g., scissors and different utensils). Of the three of them, only one acknowledges this omission and even discusses tools (11). In that study, they speculate that one possibility is that the failure of tool-selectivity to emerge from their analysis may be because their study focused on VOTC and it is often LOTC thats shows the most distinctive effects of tool selectivity. Whatever the explanation for why tool-selectivity did not emerge from the analyses performed in these studies, we believe their findings are consistent with our hypothesis. All three NSD studies visualized the loci of food-selectivity on the cortical surface, which they reported as medial and lateral to face-selectivity in LOTC and VOTC. One of these studies did a whole brain analysis and also found food-selectivity in the dorsal pathway in PCS (12). These are the same three primary regions of overlapping tool- and food-selectivity that we observed (**Fig 2**). The most notable qualitative difference was that our whole-brain results were more left lateralized. Thus, the results of these data-driven studies show that food-selectivity co-localizes, at least superficially, with regions known to be selective for tools. To clarify, we do not mean to suggest that visual cortex represents food as “the same” as tools – indeed our behavioral results speak against making such an equivalence claim (**Fig 1**). Rather, food and tool images co-localize activity in visual cortex because, like tools, food is a target of action and visual cortex may thus jointly code for tools and food for this reason – just as there is overlapping activation in LOTC for hands and tools (23) – as part of larger organization for action recognition and guidance (39,40).

### The domain-specificity of “tool-selective” visual cortex

Tools can be defined as graspable objects that extend the body during the course of actions that are used to interact with other aspects of the environment (21). An influential characterization of tool-selectivity in the visual cortex is that it reflects not specialization to represent tools as such, but rather reflects domain-specific specialization for the manipulation of objects in the service of action (41,42). Under this view, domain-specificity is a reflection of networks not specific brain areas, and specific brain regions instead represent constellation of properties related to the domain such as what objects are, how they can be manipulated, and their functional purpose. Food items can be thought of as tool-like, under the above definition, when grasped directly during the course of self-directed action. As we have shown, images of graspable food items elicit similar patterns of activation in both the ventral and dorsal visual streams, in regions that have been associated with classification of tools and their associated actions (39,40)

Our results suggest a modification to this picture of the domain-specificity associated with tool-selective brain regions. Manipulability and body extension, the hallmarks of toolness, plausibly comes in degrees. Typically, a division is drawn between tools (as defined above) on the one hand, and manipulable objects that seemingly do not extend the body during action (e.g. a smartphone). The evidence that manipulable non-tool objects are represented differently was clear in the our behavioral results but not so clearly in the neural responses measured from tool-selective ROIs, contrary to previous studies (19,27). The latter result suggests the boundary between tools and manipulable objects may not be as sharp as previously proposed, which is only further reinforced by our results for graspable food items. For example, from the perspective of action, a paintbrush dipped in paint and put to the canvas, with respect to object manipulation and body extension, is not so different from a French fry dipped in ketchup, placed in the mouth. This suggests that the texture, material, and mechanical properties that are represented for the service of action will not be well-captured by our models if we restrict ourselves to thinking of object manipulability solely in terms of artefacts like tools without considering other objects as well. This accords with a picture of visual cortex as having an action focused network that extends through ventral and dorsal visual cortex (40). Our results suggest it is this network that is engaged when viewing graspable food items.

### Comparing (neural) representations of object type and shape

Previously it has been proposed that the apparent tool-selectivity in the visual cortex may result from shape biases in stimulus selection, since many canonical tools (e.g., a hammer) are elongated, or have a higher aspect-ratio in their shape profiles, compared to other types of objects (29,30). In line with previous studies (27,31), our results do not support such a hypothesis, since we controlled for aspect-ratio in our stimulus design. Although we found a small univariate effect for shape in LOTC-tool, our RSA results found that any variance explained by the object shape RDM was shared with the object type RDM.

However, our results are discordant with other studies that have contrasted the neural representation of object type and shape in visual cortex more generally in the interest of showing that neural effects of object category cannot be accounted for by their low-level visual properties (17). These studies also used behavioral similarity ratings to generate object type and shape RDMs that were wholly orthogonal and found that each were correlated with neural RDMs for different portions of OTC and parietal cortex (43–45). Reanalysis of this data further found that layers of multiple DNNs similarly correlated with these independent object type and shape RDMs (35). Contrary to these studies, we did not find that the behavioral RDMs were orthogonal, and in fact they were strongly correlated. This is likely a result of these prior studies using multiple arrangement tasks to extract similarity ratings, which involves global judgments about arrays of stimuli at once (46). It is therefore possible that the difference in results we observe is a byproduct of the choice of similarity task. In contrast to RDMs obtained from the odd-one-out triplet task, constructing RDMs from the multiple arrangement task would not reflect fine grained comparisons with respect to object type and or shape.

### Future directions

There are many fruitful directions for further research based on our findings. First, the overlap between food and tool responses could be further explored. One prediction is that, like with tools, the neural response to food items in LOTC will also overlap with activation for hands as found in previous work (23). However, it remains to be seen what form the overlap takes at higher spatial resolution (47) or possibly at intracranial recording sites. Second, we found weaker effects in PCS-tool region. This might be because we used static images of objects without an actor being present. This region responds more strongly to depictions of actions (39,40,48). Thus, one direction of inquiry is to measure neural responses during viewing of videos of food manipulation and eating along with action types that are not food related, and measure the similarity of these actions and relate them to neural responses. Third, often item level studies of food images focus on the foods’ form of preparation or their nutritional value (9). Our behavioral results suggest the possibility of further exploring grouping food items based on the way they are acted upon and manipulated. The focus on action may also reveal differences between the dorsal and ventral pathways. In line with this, activation may vary depending on the task performed in scanner if manipulation or nutritional value is cued in separate tasks (43,49). Fourth, although we selected objects to be highly graspable and not require utensils for consumption, it remains to be seen whether other food items that typically require utensils are similarly represented. We hypothesize that manipulability is graded, and this is something that may extend to food at different stages of preparation and portioning. The present study only sampled from one extreme of this dimension. Fifth, how we engage with food is dependent on individual differences and cultural practices. For example, whether one uses utensils to eat a pizza slice is culturally determined and has associated social norms depending on context (e.g., using utensils when eating pizza in Italy, but not the USA). Sixth, there is also room to look at how food responses develop over time and with experience. For example, our preferences and food dietary restrictions may impact how we visually represent the objects, much as greater expertise influences the shared representation of hands and tools (50).

## Conclusion

How does the visual system represent complex, ecologically important stimuli like food? We conjectured that food items share many qualities in common with tools and may be represented similarity in visual cortex. In support of this we carried out an experiment that compared the location, magnitude, and pattern of neural activity for graspable food items to those elicited by tools in the visual cortex. In line with our conjecture food items produced similar neural responses to tools suggesting that they recruit a network of visual areas specialized for representing objects as focal points of action and manipulation. These results provide important insights about how the visual system digests food as an ecologically important aspect of the visible world.

## Materials and methods

### Participants

20 adult volunteers (11 women; mean, 22.9; age range, 18-27 years) participated in the fMRI experiment. For the online behavioral experiments, volunteers within the US were recruited via Amazon’s mechanical Turk: N=424 for the object type triplet task; N = 465 for the object shape triplet task; and N = 281 for the rating tasks. For the rating tasks, participants were randomly assigned to participate in one of the seven different tasks (N = 39 to 41). Online volunteers were excluded from participating in multiple experiments. All fMRI volunteers were predominantly right-handed, had normal or corrected-to-normal vision, and provided written informed consent for participation in the experiments. All online participants indicated their informed consent by button press. All experiments were approved by the institutional review board of the National Institutes of Health and all methods were performed in accordance with the relevant guidelines and regulations.

### Stimuli

The primary stimuli consisted of 48 grey scale natural images of objects without scene backgrounds (**Fig 1A**). For the fMRI experiment, all images were scaled or cropped to 800 x 800 pixels and subtended ∼10 degrees of visual angle in the scanner. For the online experiments the images, were rescaled to 400 x 400 pixels. The images were selected to include eight exemplars each of six object groupings: graspable food items (Food Items), tools used in food preparation (Food Tools), tools used in actions directed at other objects (Other Tools), tools used in self-directed action (self-Tools), manipulable non-tool artefacts (Manipulable Objects), and animal exemplars (Animals). The food items stimuli were selected to be highly familiar to American participants and stereotypically consumed using the hands alone, without the aid of utensils. The shape profiles of these images were counterbalanced across object groupings in order to have higher and lower aspect ratios, or more or less elongation, across different object orientations (44,45).

### Triplet task experiments

Two groups of participants viewed triplets of the stimuli and were given one of two instructions: (i) to judge which stimulus was unlike the other two while ignoring the shape profile of the stimuli; (ii) or to judge which had a shape unlike the other two while ignoring object type (Hebart et al. 2019). On each trial the triplet of stimuli appeared on the screen until participants made a response. Since there were 17296 unique triplets of stimuli these were randomly chunked into 412 sets of 42 trials (with one set of 34 trials). After being recruited via Amazon’s Mechanical Turk participants were randomly assigned to complete a given chunk of trials. Participants were also allowed to complete multiple chunks. Stimulus presentation and control were online via PsychoPy3 (51) hosted on pavlovia.org. Data was collected in three stages based on the following measure of quality control: if participants responded uniformly across all trials and/or had a median reaction time < 200 ms or > 10000 ms, their data were discarded (object type: 61/424 = 14% of data; object shape: 78/465 = 17% of data), and the incomplete sets were collected again until almost all sets of trials were successfully completed (object type: 396/412 = 96 % of trial sets; object shape: 391/412 = 95 %). Data were combined by counting the number of times a stimulus was the odd one out relative to another stimulus and dividing by the total number of trials the two stimuli appeared together. To determine the split-half-reliability of the matrix, the data was randomly split in half and the resulting RDMs were correlated with each other. The reported split half reliability is the mean correlation of 1000 random splits. The resulting average coefficient value was transformed using the Spearman–Brown formula. This resulting value gives an estimate of the reliability of the group-averaged data, based on the full sample size (34). To quantify the structure of the triplet RDMs, two model matrices were created by dummy coding the relative similarity of the objects based on type and shape, which were then correlated (Spearmen’s ρ) with the triplet task RDMs (**Fig 1B-C**).

### Rating experiments

Participants were tasked with rating the stimuli on a scale of 1 to 7 based on one of seven different instructions. Three were adapted from (19): how difficult would it be to pantomime the use of the object (“play charades”); how central is the form of movement of the object central to its use; and how frequently do you interact with the object with your hands? Four were adapted from (20): how much does one’s arm move when using the object; How much force is required to use the object; how much does the hand make contact when using the object; to what extent is the object used near or far from the body? Each participant completed one trial for each stimulus, which were presented in random order and the next trial did not commence until a response was collected. Stimulus presentation and control were online via Psychtoolkit (52). The split-half reliability of the ratings for each task was assessed by randomly splitting the data 1000 times and then taking the average. The resulting average coefficient value was transformed using the Spearman–Brown formula. This resulting value gives an estimate of the reliability of the group-averaged data, based on the full sample size (34). For the tasks from (19) we followed their procedure of reporting the average response for object type. To determine the relationship between the results for the different action rating tasks, the pairwise correlations between the mean ratings were calculated and PCA was carried following the approach of (20).

### Scanning procedures

The fMRI experiment consisted of a single session of eight experimental runs followed by four localizer runs and an anatomical scan. Stimuli were presented on a 32” BOLD screen (Cambridge Research Sustems Ltd, UK) placed behind the bore of the scanner and viewed through a surface mirror attached to the head coil, at a viewing distance of ∼166 cm. Stimulus presentation was controlled via a PC computer running PsychoPy3 (51).

Using a rapid event related design, each experimental run consisted of two random sequences of the 48 stimulus images. Each trial began with the stimulus centrally presented for 500 ms along with a fixation bullseye, followed by a jittered inter-stimulus interval (ISI) in which the bullseye continued to be presented. ISI jitter was achieved by adding different numbers of intervals of 50 ms to a base ISI of 1500 ms from a set pool of intervals, ensuring that the mean ISI was 2000 ms and the total duration was identical across runs. On each trial participants indicated whether the object in the image was larger or smaller than their hand by responding with the index and middle fingers of their right hand. The mapping between response options (larger/smaller) and response buttons (left/right) was counterbalanced across runs. Experimental runs had a total duration of 4 min 20 s.

For the localizer runs a block design was used with five stimulus types: animals, foods, objects (furniture and vehicles), tools, and box-scrambled versions of the object images, with 18 images of each stimulus type. Like with the main experimental stimuli, the food images were selected to be of food items that are directly grasped when being consumed and to be familiar to American volunteers. Each image in a block appeared for 400 ms followed by 400ms fixation with four repeats of each stimulus block type per run. All five series of image types were presented sequentially in each stimulus block in pseudorandom order, followed by a 16 s fixation period. Localizer runs had a duration of 6 min 32 s. To maintain their attention during a run, participants indicated when one of the images was repeated later in an image series, which occurred once each for two different randomly selected images per image type per block. A fixation bull’s-eye was centrally presented continuously throughout each run.

### Acquisition parameters

Data acquisition was conducted using a 3T Siemans Prisma scanner, with a 32-channel coil, at the Clinical Center at the National Institutes of Health in Bethesda, USA. Functional MRI volumes were acquired using a two-dimensional (2D) multiband (MB) T2*-weighted echo planar imaging sequence with GRAPPA. MB = 2; repetition time, 2000ms; echo time, 28 ms; flip angle, 65 °; field of view, 212; voxel size = 2 x 2 x 2 mm; matrix size = 106 x 106. Each volume consisted of 54 axial slices (no gap) aligned to encompass as much of the cortex as possible and included all of the occipital and temporal lobes. Typically, this position resulted in the exclusion of the most superior portions of the parietal and frontal lobes from the volume. The T1-weighted anatomic volumes were acquired for each participant using an MPRAGE sequence, 1 x 1 x 1 mm resolution.

### fMRI preprocessing and analysis

First, prior to any analysis functional images were converted from dicom to AFNI format and any personal identifying information was removed from the image header files. Anatomical volumes were converted from dicom to NIFTI format, the face was removed using pydeface (https://github.com/poldracklab/pydeface) and personal identifying information was removed from the image header files. Second, freesurfer was then used for surface reconstruction from the anatomical images (53) and then SUMA was used to translate the freesurfer outputs into a format suitable for AFNI (54). Third, for each experimental and localizer run, the following preprocessing steps were carried out using AFNI (55): the first four acquisitions were removed from the time series; despiking was used to smooth the time series; the volumes were motion corrected by aligning them to the minimal outlier in the time series; the volumes were aligned to the individual anatomical image; and the functional volumes were masked based on the anatomical image. Smoothing (4 mm FWHM) was carried out only for the localizer runs. Fourth, after preprocessing, the time series for the experimental and localizer runs were separately modeled using GLMs with the following settings, with regressors for each stimulus event and block type: AFNI’s SPMG1 was used as the basis function; motion correction parameters were included as predictors; generalized least squares times fit was calculated based on the residual maximum likelihood estimation (REML); acquisitions were censured that had excessive motion or excessive outliers. This resulted in a single beta estimate for each condition from across the runs.

### Group level analysis

To carry out the group analysis the results of the REML GLM were normalized into a common Talairach space and then mixed-effects multilevel analysis (MEMA) was carried out based on the (26). MEMA takes into account both variation across participants and the quality of effect estimates of each condition based on the beta values and t-statistics of the regression models for each participant. MEMA was carried out for three functional contrasts, tool > animal, food > animal, and object > animal. Results were thresholded at p < 0.05, uncorrected. The results were visualized using SUMA (54).

### Overlap analysis

Anatomical masks were defined for the left hemisphere based on a conjunction of parcels from Human Connectome Project (56). These conjunctions of surface parcels were then converted from MNI space to the native space of individual participants. The overlap for the two contrasts was carried out in two ways within. First, a liberal threshold of 0.05 using a cluster value of 40 was used to define significant contrast maps within the individual participant space. The overlap between these two results was then determined. For this one participant was excluded for LOTC and one for VOTC, all of this in the left hemisphere. The Jaccard index, or the intersection as a proportion of the union of two sets of binary values, was used as the measure of overlap. Where F and T are the voxels of the food- and tool-selective maps above threshold, the index is F ∩ T / F ⋃ T (28). The non-overlapping portions of the maps were also calculated by dividing F and T by F ⋃ T. Statistical significance was determined by two-sided t-tests comparing the non-overlapping portion of the food-selective map to the overlap and non-overlapping portion of the tool-selective map. Second, the topmost selective voxels, without clustering, were selected within the masked area in steps of 50 voxels (50 to 500). Again, at each stage the proportion overlap was calculated based on Jaccard index, or the intersection of the top responding voxels for each contrast proportional to their union.

### Defining regions of interest

Five ROIs were defined in the following way from the results of the localizer GLM. Using the same anatomical masks as those used for calculating contrast overlap, maps were clustered based on minimum 40 voxels (including face and edgewise neighbors), at multiple thresholds (0.001, 0.005, 0.01, 0.05). Starting with the most stringent, this was convolved with the mask and if the number of voxels was less than the minimum cluster number, a lower threshold was used. If no voxels survived the lowest threshold no ROI was created. For PCS-tool, LOTC-animal, and VOTC-animal an ROI image was created for all 20 participants, while for only 19/20 and 18/20 for LOTC-tool and VOTC-tool, respectively. The left EVC ROI consisted of voxels within the V1 parcel of LH HCP that responded to the scrambled image condition, p < 0.001, cluster threshold of 40 voxels (including face and edgewise neighbors). All resulting ROI images were converted to NIFTI for further analysis.

### Univariate analysis

The bold response across ROI voxels was averaged at the level of individual exemplars for visualization or object and shape types for statistical analysis. Statistical significance was determined by two-sided t-tests comparing the average magnitude of the responses for food items relative to other object types. To determine whether the magnitude of response was predicted by the mean values for individual exemplars, the mean responses of the rating tasks (or PCs) were correlated (Pearson’s *r*) with the individual magnitude responses in the relevant ROIs. Analysis was carried out in MATLAB using custom code and CoSMoMVPA (57).

### Representational similarity analysis

RSA was used to compare the activity patterns from the different ROIs to the triplet task data, a shape model, and layers of a suite of DNNs (58). For each comparison, representational dissimilarity matrices (RDMs) were constructed, which are matrices that are symmetrical around the diagonal and reflect the pairwise dissimilarities among all stimulus conditions. RDMs from different data modalities can be directly compared in order to evaluate the second order isomorphisms of the dissimilarities between conditions. RSA was conducted using CoSMoMVPA and custom Matlab code (57). Neural RDMs for the different ROIs for each participant were constructed using the 1-r correlation distance (58). To assess the between-participant reliability of the RDMs for each ROI, the RDM of one participant was left out and those of the remaining participants were averaged, and Pearson’s r correlated with the left-out participant’s RDM. This was conducted for all participants, and the resulting coefficients were averaged. The resulting region-specific average value was used as an estimate of the noise ceiling when correlating individual neural RDMs for each ROI with the RDMs from the other data modalities. Visualization of group-averaged neural RDMs included multidimensional scaling (MDS) with stress 1 as the criterion. The dissimilarity matrix for the object type and object shape triplet tasks were used as model RDMs. For CORnet (described below), layer-specific RDMs were constructed based on the 1–r pairwise Pearson’s correlation between the vectors of unit responses for each image. To compare RDMs from different data modalities, the bottom half of each matrix was converted to a vector, and the Spearman rank-order correlation was calculated between matrices. The median correlations across participants were tested for significance using two-sided t-tests.

### CORnet

CORnet is a family of recurrent DNN architectures with four layers that are premapped onto the areas of the ventral visual pathway in the primate brain, V1, V2, V4, and IT, along with a final decoding layer (36). We used CORnet-S, which combines skip connections with within-area recurrent connections and previously performed best “overall” on the Brain-Score benchmark (37).

We selected this architecture because it was designed to match neural responses brain regions like OTC. We used the implementation of CORnet-S found pre trained on the ImageNet stimulus set (59) as implemented in THINGsvision (60).

### Commonality analysis

Commonality analysis, or “variance partitioning” is a method for determining whether multiple predictors uniquely or jointly explain variance in the dependent variable (61) and was used in conjunction with RSA (38,62). In the present case of three predictors (a, b, c) for some dependent variable y, there were seven coefficients of determination (R^2^) for all possible combinations of predictors in a linear regression model: R^2^ _y · a_, R^2^ _y · b_, R^2^ _y · c_, R^2^ _y · ab_, R^2^ _y · ac_, R^2^ _y · bc_, R^2^ _y · abc_. The last of these is the full model, for which the variance is partitioned based on differential weighting of the R^2^ of the different models. When there are only three predictors, the partitioning can be performed using a simple weighting table, in which the vector of coefficients is multiplied with row vectors of weights for each of the unique and common variance components (63). These results were visualized with EulerAPE (64).

To carry out the multiple regression necessary for commonality analysis, RDMs were converted to vectors, and the group-averaged neural dissimilarity values were regressed on the different model or behavioral dissimilarity vectors. Significance was determined using a permutation test. For each individual participant, the rows of the bottom half of their neural RDM were independently randomly shuffled, and the resulting random vector was averaged across participants and then fit with the full model. This procedure was conducted 1000 times for each application of multiple regression. The resulting proportion of R^2^–values greater than that observed for the full model (fit to the unshuffled dissimilarities) provides the p value for the test.

## Declarations

### Author contributions

J.B.R. designed the experiment with input from M.V-P and C.I.B; J.B.R and S.A collected the data; J.B.R analyzed the data; J.B.R. wrote the manuscript with input from S.A., M.V-P and C.I.B.

### Data availability

Stimuli and raw and processed data are available at https://osf.io/n2amw/.

### Ethical approval

This research was approved by the IRB of the National Institutes of Health.

## References

1. Bracci S, Op de Beeck HP. Understanding Human Object Vision: A Picture Is Worth a Thousand Representations. Annu Rev Psychol. 2023 Jan 18;74:113–35.

2. Grill-Spector K, Weiner KS. The functional architecture of the ventral temporal cortex and its role in categorization. Nat Rev Neurosci. 2014 Aug;15(8):536–48.

3. Malcolm GL, Groen IIA, Baker CI. Making Sense of Real-World Scenes. Trends in Cognitive Sciences. 2016 Nov 1;20(11):843–56.

4. Peelen MV, Downing PE. Category selectivity in human visual cortex: Beyond visual object recognition. Neuropsychologia. 2017 Oct;105:177–83.

5. Simmons WK, Martin A, Barsalou LW. Pictures of Appetizing Foods Activate Gustatory Cortices for Taste and Reward. Cerebral Cortex. 2005 Oct 1;15(10):1602–8.

6. van der Laan LN, de Ridder DTD, Viergever MA, Smeets PAM. The first taste is always with the eyes: A meta-analysis on the neural correlates of processing visual food cues. NeuroImage. 2011 Mar 1;55(1):296–303.

7. Avery JA, Liu AG, Ingeholm JE, Gotts SJ, Martin A. Viewing images of foods evokes taste quality-specific activity in gustatory insular cortex. Proceedings of the National Academy of Sciences. 2021 Jan 12;118(2):e2010932118.

8. Avery JA, Carrington M, Martin A. A common neural code for representing imagined and inferred tastes. Progress in Neurobiology. 2023 Apr 1;223:102423.

9. Coricelli C, Toepel U, Notter ML, Murray MM, Rumiati RI. Distinct brain representations of processed and unprocessed foods. European Journal of Neuroscience. 2019;50(8):3389–401.

10. Jain N, Wang A, Henderson MM, Lin R, Prince JS, Tarr MJ, et al. Selectivity for food in human ventral visual cortex. Commun Biol. 2023 Feb 1;6(1):175.

11. Khosla M, Ratan Murty NA, Kanwisher N. A highly selective response to food in human visual cortex revealed by hypothesis-free voxel decomposition. Current Biology. 2022 Oct 10;32(19):4159–4171.e9.

12. Pennock IML, Racey C, Allen EJ, Wu Y, Naselaris T, Kay KN, et al. Color-biased regions in the ventral visual pathway are food selective. Curr Biol. 2023 Jan 9;33(1):134–146.e4.

13. Allen EJ, St-Yves G, Wu Y, Breedlove JL, Prince JS, Dowdle LT, et al. A massive 7T fMRI dataset to bridge cognitive neuroscience and artificial intelligence. Nat Neurosci. 2022 Jan;25(1):116–26.

14. Bannert MM, Bartels A. Visual cortex: Big data analysis uncovers food specificity. Current Biology. 2022 Oct 10;32(19):R1012–5.

15. Simmons WK, Rapuano KM, Kallman SJ, Ingeholm JE, Miller B, Gotts SJ, et al. Category-specific integration of homeostatic signals in caudal but not rostral human insula. Nat Neurosci. 2013 Nov;16(11):1551–2.

16. LaBar KS, Gitelman DR, Parrish TB, Kim YH, Nobre AC, Mesulam MM. Hunger selectively modulates corticolimbic activation to food stimuli in humans. Behavioral Neuroscience. 2001;115(2):493–500.

17. Bracci S, Ritchie JB, de Beeck HO. On the partnership between neural representations of object categories and visual features in the ventral visual pathway. Neuropsychologia. 2017 Oct 1;105:153–64.

18. Hebart MN, Dickter AH, Kidder A, Kwok WY, Corriveau A, Wicklin CV, et al. THINGS: A database of 1,854 object concepts and more than 26,000 naturalistic object images. PLOS ONE. 2019 Oct 15;14(10):e0223792.

19. Mahon BZ, Milleville SC, Negri GAL, Rumiati RI, Caramazza A, Martin A. Action-Related Properties Shape Object Representations in the Ventral Stream. Neuron. 2007 Aug 2;55(3):507–20.

20. Watson CE, Buxbaum LJ. Uncovering the architecture of action semantics. Journal of Experimental Psychology: Human Perception and Performance. 2014;40(5):1832–48.

21. Mahon BZ. The Representation of Tools in the Human Brain. 2020 May 12 [cited 2023 Sep 27]; Available from: https://direct.mit.edu/books/edited-volume/5456/chapter/3967027/The-Representation-of-Tools-in-the-Human-Brain

22. Beauchamp MS, Lee KE, Haxby JV, Martin A. fMRI Responses to Video and Point-Light Displays of Moving Humans and Manipulable Objects. Journal of Cognitive Neuroscience. 2003 Oct 1;15(7):991–1001.

23. Bracci S, Cavina-Pratesi C, Ietswaart M, Caramazza A, Peelen MV. Closely overlapping responses to tools and hands in left lateral occipitotemporal cortex. J Neurophysiol. 2012 Mar 1;107(5):1443–56.

24. Chao LL, Haxby JV, Martin A. Attribute-based neural substrates in temporal cortex for perceiving and knowing about objects. Nat Neurosci. 1999 Oct;2(10):913–9.

25. Valyear KF, Cavina-Pratesi C, Stiglick AJ, Culham JC. Does tool-related fMRI activity within the intraparietal sulcus reflect the plan to grasp? NeuroImage. 2007 Jan 1;36:T94– 108.

26. Chen G, Saad ZS, Nath AR, Beauchamp MS, Cox RW. FMRI group analysis combining effect estimates and their variances. NeuroImage. 2012 Mar 1;60(1):747–65.

27. Chen J, Snow JC, Culham JC, Goodale MA. What Role Does “Elongation” Play in “Tool-Specific” Activation and Connectivity in the Dorsal and Ventral Visual Streams? Cerebral Cortex. 2018 Apr 1;28(4):1117–31.

28. Kung CC, Peissig JJ, Tarr MJ. Is Region-of-Interest Overlap Comparison a Reliable Measure of Category Specificity? Journal of Cognitive Neuroscience. 2007 Dec 1;19(12):2019–34.

29. Almeida J, Mahon BZ, Zapater-Raberov V, Dziuba A, Cabaço T, Marques JF, et al. Grasping with the eyes: The role of elongation in visual recognition of manipulable objects. Cogn Affect Behav Neurosci. 2014 Mar 1;14(1):319–35.

30. Sakuraba S, Sakai S, Yamanaka M, Yokosawa K, Hirayama K. Does the human dorsal stream really process a category for tools? Journal of Neuroscience. 2012;32(11):3949–53.

31. Matić K, Op de Beeck H, Bracci S. It’s not all about looks: The role of object shape in parietal representations of manual tools. Cortex. 2020 Dec 1;133:358–70.

32. Kriegeskorte N. Deep Neural Networks: A New Framework for Modeling Biological Vision and Brain Information Processing. Annu Rev Vis Sci. 2015 Nov 24;1(1):417–46.

33. Lindsay GW. Convolutional Neural Networks as a Model of the Visual System: Past, Present, and Future. Journal of Cognitive Neuroscience. 2021 Sep 1;33(10):2017–31.

34. Ritchie JB, Zeman AA, Bosmans J, Sun S, Verhaegen K, Beeck HPO de. Untangling the Animacy Organization of Occipitotemporal Cortex. J Neurosci. 2021 Aug 18;41(33):7103– 19.

35. Zeman AA, Ritchie JB, Bracci S, Op de Beeck H. Orthogonal Representations of Object Shape and Category in Deep Convolutional Neural Networks and Human Visual Cortex. Sci Rep. 2020 Feb 12;10(1):2453.

36. Kubilius J, Schrimpf M, Kar K, Rajalingham R, Hong H, Majaj N, et al. Brain-Like Object Recognition with High-Performing Shallow Recurrent ANNs. In: Advances in Neural Information Processing Systems [Internet]. Curran Associates, Inc.; 2019 [cited 2023 Sep 28]. Available from: https://proceedings.neurips.cc/paper/2019/hash/7813d1590d28a7dd372ad54b5d29d033-Abstract.html

37. Schrimpf M, Kubilius J, Lee MJ, Ratan Murty NA, Ajemian R, DiCarlo JJ. Integrative Benchmarking to Advance Neurally Mechanistic Models of Human Intelligence. Neuron. 2020 Nov 11;108(3):413–23.

38. Groen II, Greene MR, Baldassano C, Fei-Fei L, Beck DM, Baker CI. Distinct contributions of functional and deep neural network features to representational similarity of scenes in human brain and behavior. Tsao DY, editor. eLife. 2018 Mar 7;7:e32962.

39. Orban GA, Caruana F. The neural basis of human tool use. Frontiers in Psychology [Internet]. 2014 [cited 2024 Jan 5];5. Available from: https://www.frontiersin.org/articles/10.3389/fpsyg.2014.00310

40. Wurm MF, Caramazza A. Two ‘what’ pathways for action and object recognition. Trends in Cognitive Sciences. 2022 Feb 1;26(2):103–16.

41. Mahon BZ. Chapter 13 - Domain-specific connectivity drives the organization of object knowledge in the brain. In: Miceli G, Bartolomeo P, Navarro V, editors. Handbook of Clinical Neurology [Internet]. Elsevier; 2022 [cited 2023 Jan 12]. p. 221–44. (The Temporal Lobe; vol. 187). Available from: https://www.sciencedirect.com/science/article/pii/B9780128234938000286

42. Mahon BZ, Caramazza A. What drives the organization of object knowledge in the brain? Trends in Cognitive Sciences. 2011 Mar 1;15(3):97–103.

43. Bracci S, Daniels N, Op de Beeck H. Task Context Overrules Object- and Category-Related Representational Content in the Human Parietal Cortex. Cerebral Cortex. 2017 Jan 1;27(1):310–21.

44. Bracci S, Op de Beeck H. Dissociations and Associations between Shape and Category Representations in the Two Visual Pathways. J Neurosci. 2016 Jan 13;36(2):432–44.

45. Ritchie JB, de Beeck HO. Using neural distance to predict reaction time for categorizing the animacy, shape, and abstract properties of objects. Sci Rep. 2019 Sep 13;9(1):13201.

46. Kriegeskorte N, Mur M. Inverse MDS: Inferring Dissimilarity Structure from Multiple Item Arrangements. Frontiers in Psychology [Internet]. 2012 [cited 2024 Jan 5];3. Available from: https://www.frontiersin.org/articles/10.3389/fpsyg.2012.00245

47. Schwarzlose RF, Baker CI, Kanwisher N. Separate Face and Body Selectivity on the Fusiform Gyrus. J Neurosci. 2005 Nov 23;25(47):11055–9.

48. Peeters R, Simone L, Nelissen K, Fabbri-Destro M, Vanduffel W, Rizzolatti G, et al. The Representation of Tool Use in Humans and Monkeys: Common and Uniquely Human Features. J Neurosci. 2009 Sep 16;29(37):11523–39.

49. Vaziri-Pashkam M, Xu Y. Goal-Directed Visual Processing Differentially Impacts Human Ventral and Dorsal Visual Representations. J Neurosci. 2017 Sep 6;37(36):8767–82.

50. Schone HR, Maimon-Mor RO, Baker CI, Makin TR. Expert Tool Users Show Increased Differentiation between Visual Representations of Hands and Tools. J Neurosci. 2021 Mar 31;41(13):2980–9.

51. Peirce J, Gray JR, Simpson S, MacAskill M, Höchenberger R, Sogo H, et al. PsychoPy2: Experiments in behavior made easy. Behav Res. 2019 Feb 1;51(1):195–203.

52. Stoet G. PsyToolkit: A Novel Web-Based Method for Running Online Questionnaires and Reaction-Time Experiments. Teaching of Psychology. 2017 Jan 1;44(1):24–31.

53. Fischl B. FreeSurfer. NeuroImage. 2012 Aug 15;62(2):774–81.

54. Saad ZS, Reynolds RC, Argall B, Japee S, Cox RW. SUMA: an interface for surface-based intra- and inter-subject analysis with AFNI. In: 2004 2nd IEEE International Symposium on Biomedical Imaging: Nano to Macro (IEEE Cat No 04EX821) [Internet]. 2004 [cited 2023 Oct 3]. p. 1510–1513 Vol. 2. Available from: https://ieeexplore.ieee.org/abstract/document/1398837

55. Cox RW. AFNI: Software for Analysis and Visualization of Functional Magnetic Resonance Neuroimages. Computers and Biomedical Research. 1996 Jun 1;29(3):162–73.

56. Glasser MF, Coalson TS, Robinson EC, Hacker CD, Harwell J, Yacoub E, et al. A multi-modal parcellation of human cerebral cortex. Nature. 2016 Aug;536(7615):171–8.

57. Oosterhof NN, Connolly AC, Haxby JV. CoSMoMVPA: Multi-Modal Multivariate Pattern Analysis of Neuroimaging Data in Matlab/GNU Octave. Frontiers in Neuroinformatics [Internet]. 2016 [cited 2023 Oct 3];10. Available from: https://www.frontiersin.org/articles/10.3389/fninf.2016.00027

58. Kriegeskorte N, Mur M, Bandettini P. Representational similarity analysis - connecting the branches of systems neuroscience. Frontiers in Systems Neuroscience [Internet]. 2008 [cited 2023 Feb 21];2. Available from: https://www.frontiersin.org/articles/10.3389/neuro.06.004.2008

59. Deng J, Dong W, Socher R, Li LJ, Li K, Fei-Fei L. ImageNet: A large-scale hierarchical image database. In: 2009 IEEE Conference on Computer Vision and Pattern Recognition [Internet]. 2009 [cited 2023 Oct 3]. p. 248–55. Available from: https://ieeexplore.ieee.org/abstract/document/5206848

60. Muttenthaler L, Hebart MN. THINGSvision: A Python Toolbox for Streamlining the Extraction of Activations From Deep Neural Networks. Frontiers in Neuroinformatics [Internet]. 2021 [cited 2023 Oct 3];15. Available from: https://www.frontiersin.org/articles/10.3389/fninf.2021.679838

61. Seibold DR, McPhee RD. Commonality Analysis: A Method for Decomposing Explained Variance in Multiple Regression Analyses. Human Communication Research. 1979 Jun 1;5(4):355–65.

62. Lescroart MD, Stansbury DE, Gallant JL. Fourier power, subjective distance, and object categories all provide plausible models of BOLD responses in scene-selective visual areas. Frontiers in Computational Neuroscience [Internet]. 2015 [cited 2023 Oct 18];9. Available from: https://www.frontiersin.org/articles/10.3389/fncom.2015.00135

63. Nimon K, Reio TG. Regression Commonality Analysis: A Technique for Quantitative Theory Building. Human Resource Development Review. 2011 Sep 1;10(3):329–40.

64. Micallef L, Rodgers P. eulerAPE: Drawing Area-Proportional 3-Venn Diagrams Using Ellipses. PLOS ONE. 2014 Jul 17;9(7):e101717.

